# Simulation of multiple microenvironments shows a putative role of RPTPs on the control of Epithelial-to-Mesenchymal Transition

**DOI:** 10.1101/609263

**Authors:** Ricardo Jorge Pais

## Abstract

Epithelial-to-Mesenchymal Transition (EMT) together with Mesenchymal-to-Epithelial Transition (MET) are two natural processes thought to participate in the process of metastasis. Multiple signals from the microenvironment have been reported to drive EMT. However, microenvironment signals that control EMT and promote MET are still unknown. Here, we analysed a regulatory network of EMT involving 8 microenvironment signals to evaluate the role of cell contact signals on the switching between Epithelial and Mesenchymal-like phenotypes. The results demonstrated that RPTP activation by cell contacts have the potential to control EMT and promote MET in the presence of 5 EMT driving signals under physiological scenarios. These simulations also showed that hypoxia inhibits the RPTPs capacity of controlling EMT. In this case, FAT4 activation by cell contacts functions as an alternative control mechanism of EMT except under chronic inflammation, providing a theoretical explanation for the observed correlation between hypoxia and metastasis under chronic inflammation. Taken together, we propose here a natural control mechanism of EMT that supports the idea that this process is tightly regulated by the microenvironment.

**Highlights:**

- Cell contact dependent RPTP inhibit EMT and triggers MET in the presence of 5 EMT driving signals *in silico*.
- A proposed molecular mechanism for the control of EMT by cell contact dependent RPTPs.
- A proposed explanation for the observed MET *in vitro* and the correlation between hypoxia and metastasis *in vivo*.

## 1. Introduction

EMT is a complex and reversible trans-differentiation process observed in embryogenesis and wound healing, where a cell loses the cell-cell adhesion to its neighbours and gains single cell migration capacity through the ECM [1–3]. In cancer, the transitions between Epithelial-like and Mesenchymal-like phenotypes are believed to participate in the process of metastasis of carcinomas mainly through driving cancer migration the and respective colonization in secondary sites by MET [1,4]. The microenvironment is considered to be the main factor that controls the transitions between these phenotypes [5,6]. Multiple signals from the tumour microenvironment such as ECM stiffness, inflammatory signals, hypoxia, growth factors and Delta-Notch have already been demonstrated *in vitro* to promote EMT in cancer cells [7–10]. However, microenvironment signals that prevent EMT and promote MET are not yet reported [11,12]. This has been a huge challenge in cancer research due to the multiplicity of existing signals coming from the ECM and neighbouring cells. Thus, finding good candidates as critical signals that control the interchange between migrating (Mesenchymal-like) and non-migrating cancer cells (Epithelial-like) would provide an advance in cancer research.

The hypothesis that cell contact signals from neighbouring cells in the microenvironment affect cell fate has been considered in many biological processes [10,13]. Receptor Protein Tyrosine Phosphatases (RPTP) and FAT4 are two membrane proteins receptors that are strongly activated by cell contacts and reported as capable of inhibiting the growth factor signalling (RPTP) and the Wnt signalling (FAT4), two pathways involved in EMT [12,14,15]. Thus, we hypothesise that RPTP and/or FAT4 may be capable of controlling EMT in the presence of multiple signals that stimulate EMT.

Logical modelling of regulatory networks have been successfully applied for exploring multiple hypotheses, describing observed behaviours and identifying novel biomarkers in cancer [16–19]. To analyse our hypothesis, we used a logical network model of the regulation of two critical cell adhesion properties involved in EMT, which accounts for a total of 8 key microenvironment signals [20]. This model is so far the most complete model in terms on key pathways in EMT and includes the regulatory effects of RPTP and FAT4 activation by cell contacts. Here, we report the main results of this model that evaluates our hypothesis under several physiological scenarios and propose a natural control mechanism of EMT.

## 2. Methods

### 2.1 Modelling framework

The mathematical framework used in this work was the logical formalism, initially proposed by René Thomas [21]. This approach consists in defining an interaction map that reflects the regulatory network involved, which contains the regulators (nodes) connected through arcs representing the activations or inhibitions (interactions). In this framework, the nodes are the variables, which can be binary (Boolean) or multi-valued, describing discrete qualitative states of biological activity/concentration for the respective network components (strong, intermediate or high). The behaviours of the model are defined by logical functions, which result in the evolution of the variables towards attractors, according to an update scheme. Here, we are focused on a type of attractors called stable states (also called point attractors), which are fixed points where functions are no longer updatable.

### 2.2 Network model

Modelling the transitions between phenotypes was performed using a literature-based logical model for the regulation of two critical cell adhesion properties in EMT previously described in [20]. This model was developed accounting for the published regulatory network models of EMT and further extended to include cell-cell contact dependent RPTP and FAT4 in the microenvironment [16,17]. Logical functions of this model were developed to abstract the biological mechanisms involved in the activation of each model component (e.g. translation and phosphorylation). Briefly, this model is composed by a total of 51 regulatory components (nodes) and 134 regulatory interactions, accounting for TGFβ, Integrin, Wnt, AKT, MAPK, HIF1, Notch, and Hippo signalling (Figure 1A). The model also considers the transcriptional regulation of E-cadherin and the post-transcriptional regulation of E-cadherin/β-catenin/p120 complex. The model inputs include 3 EMT relevant inflammatory signals (IL6, ROS, and the ECM stiffness), 2 growth factors (EGF and HGF) and 3 cell-cell contact signals (DELTA, FAT4L, and RPTPL). These nodes were associated with Boolean variables defining basal (value 0) or high (value 1) degrees of activity. The model outputs were associated with phenotypes (Figure 1B) according to their typical cell adhesion properties [22,23]. For further details on model components and interactions see supplementary file CAmodeldoc.xhtml, publicly available on https://github.com/rjpais/CALMproj. Here, a copy of the model (CAmodel.zginml) can be downloaded. Moreover, this model was extensively validated by comparing its results against multiple phenotypic and activity observations from published experiments on Epithelial-like cell lines (see Table S1 and Figure S2 in supplementary data).

**Figure 1.**
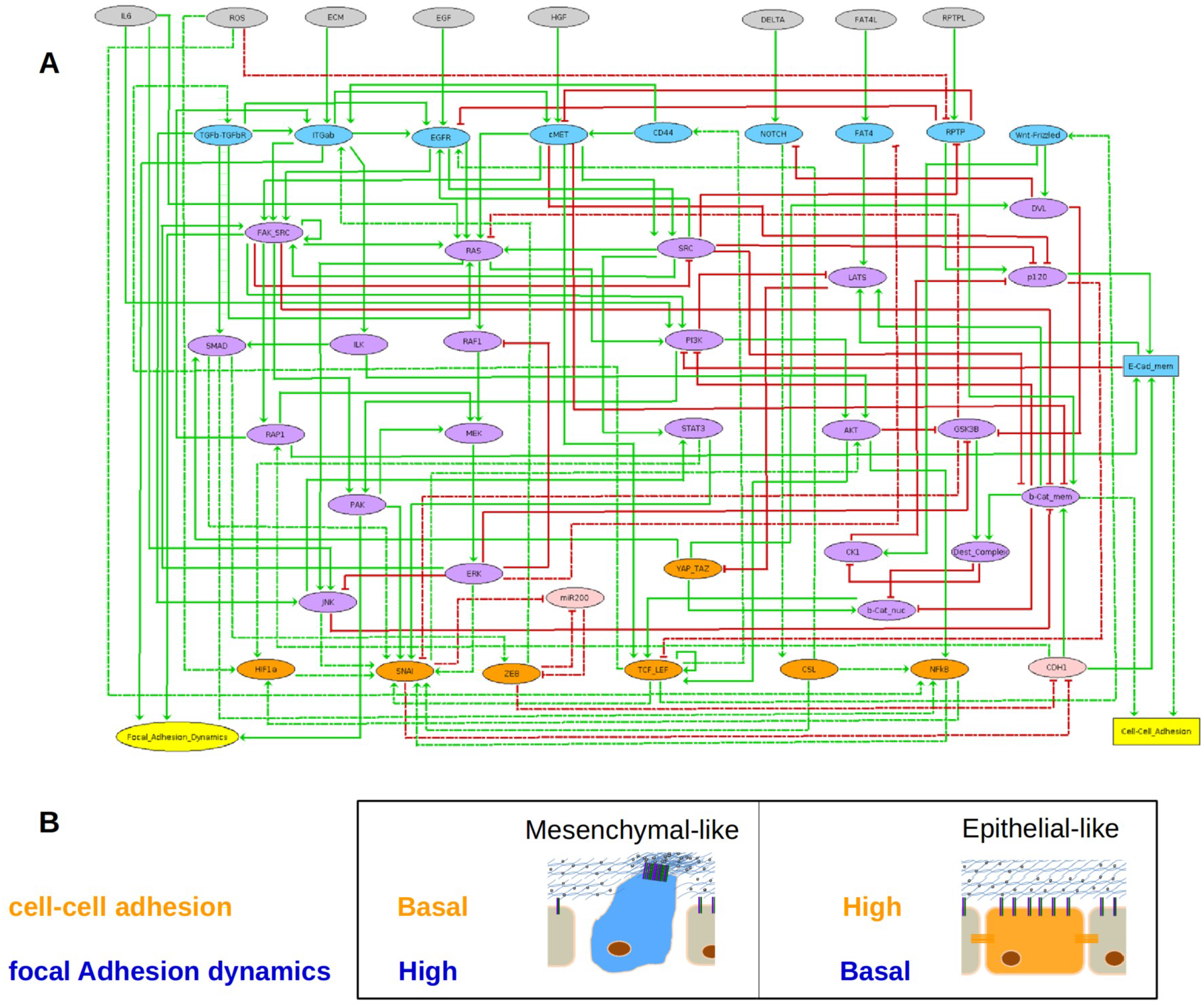
Network model (A) and related phenotypes (B). Microenvironment signals are depicted in grey, membrane protein/receptors in blue, signalling components in purple, transcription factors in orange, genes and RNA in pink, cell adhesion properties in yellow; elliptic nodes denote Boolean variables and rectangular nodes denote multi-valued variables; red arcs are inhibitions and green arcs activations. Fast interactions are denoted by plain arcs and slow interactions by dashed arcs.

### 2.3 Simulations of physiological scenarios

Simulations of physiological scenarios with the model were carried using GINsim, a free software tool for modelling regulatory networks [24]. The reachable stable states and their associated phenotypes were obtained in GINsim by generating the state transition graph with the method described in [24,25]. Model inputs were set in each simulation to mimic the cell microenvironment associated with a particular physiological scenario, starting from a defined stable state associated to Epithelial-like or Mesenchymal-like phenotype (see Figure S1 in supplementary data file). All simulations were run under asynchronous updating policy, according to rules of priority that accounted for timescale constraints using the method described in [25,26]. Knockout perturbations on regulatory interactions were also defined in GINsim by setting the values of the effects of regulators to 0 and analysed through simulation of a particular physiological scenario.

## 3 Results

Simulation of relevant physiological variations that epithelial tissues may be exposed allowed us to explore the role of cell contact dependent activation of RPTP ligands (RPTPL), FAT4 ligands (FAT4L) on the control of EMT. The outcomes from simulations resulted in predicted microenvironmental variations for the transitions between Epithelial-like and Mesenchymal-like phenotypes (Figure 2). The results showed that EMT was not compatible with high degree of RPTPL signal in the microenvironment under conditions that represent tissue growth, chronic inflammation and healthy epithelia with high activity of DELTA. Similar incompatibility was also obtained for FAT4L, except for the case of chronic inflammation. Interestingly, the model predicted that MET can only be triggered under high RPTP activity (RPTPL = 1). Simulations also showed that MET was achieved using microenvironment inputs compatible with tissue growth, chronic inflammation, DELTA signal and healthy epithelia conditions. This indicates that RPTP activation triggers MET in the presence of EMT inducing signals such as EGF, HGF, ECM stiffness and DELTA. These results suggest cell contact dependent RPTP can be a key driver of MET, explaining the observed MET in cancer cell lines due to signals from cocultured normal Epithelial cells [27]. Further, simulations showed that the model input conditions that mimic hypoxia in combination with tissue growth, chronic inflammation or DELTA signal were capable of inhibiting the RPTP control over EMT and preventing MET. This is explained by the oxidative inhibition of RPTP by ROS generated under hypoxia conditions, which in turn was accounted in the model as a regulatory interaction [28,29]. On the other hand, hypoxia could not inhibit FAT4L capacity to prevent EMT, which suggests that it could only have an effect under chronic inflammation conditions.

**Figure 2.**
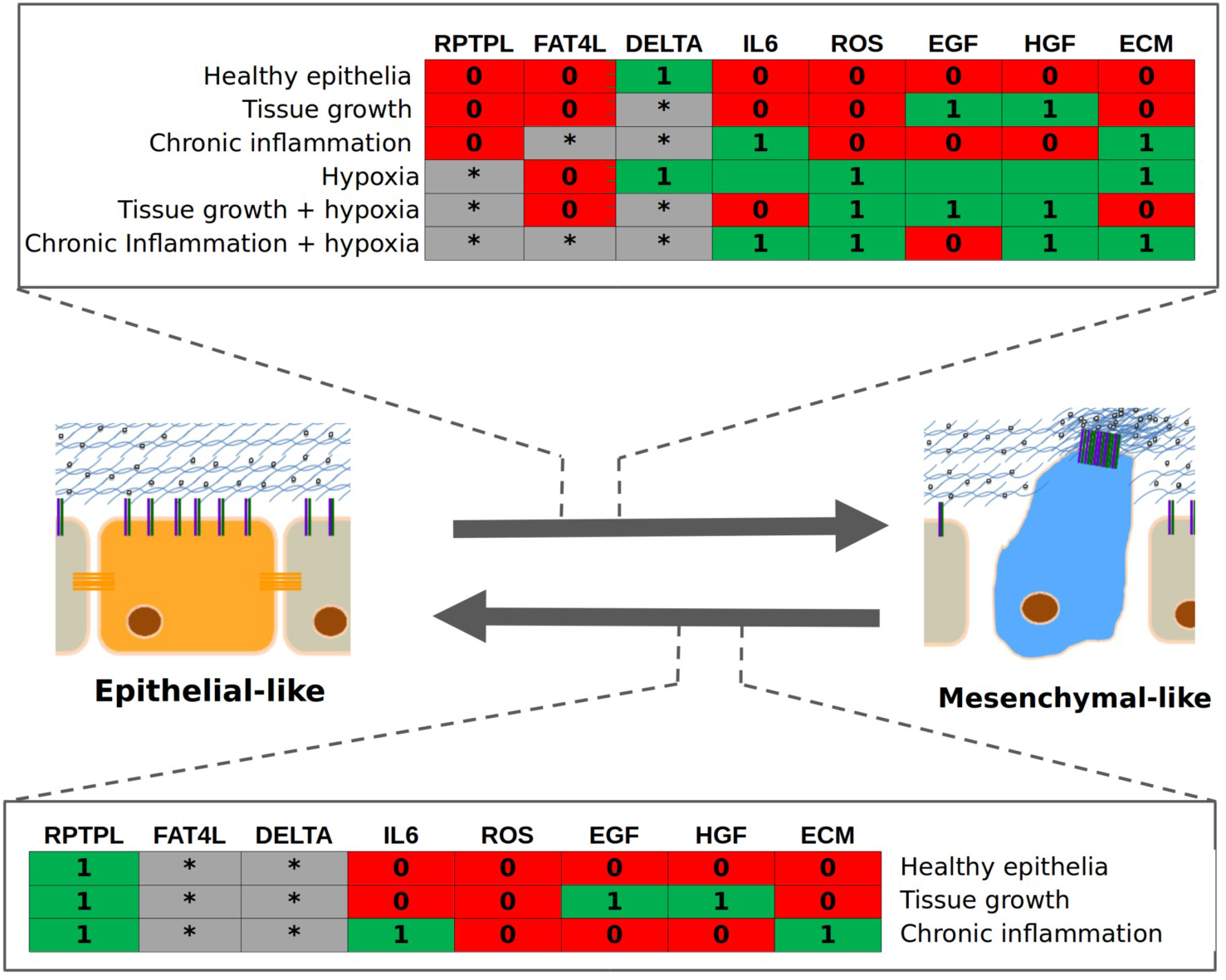
Simulation of the transitions between model phenotypes under physiological scenarios. Transitions between phenotypes are represented by dark grey arrows and the compatible conditions in linked tables. In tables, green(1) denotes high activity, red(0) denotes basal activity and grey(*) denotes all possible degrees. Details and references on the input choice for physiological scenarios indicated in table S2 and the initial stable states of phenotypes are presented in supplementary data file.

To understand the mechanism by which RPTP control of EMT and promote MET, we analyzed the impact of knockouts of RPTP regulatory effects included in the network model. For this purpose, we simulated EMT under a combination of multiple EMT driving signals and analyzed the resulting model outputs (Table 1). The results demonstrated that removing the RPTP regulatory interactions with EGFR and cMET allowed the formation of stable phenotypes with Mesenchymal-like migration properties. This indicates that these molecular interactions are required for the control of EMT. The results also showed that the knockout of the regulatory interaction between RPTP and p120 only destabilized Epithelial-like adhesion properties, showing that this interaction is also important for the control of EMT.

**Table 1.** Effects of the KO of regulatory interactions between RPTP and its targets on the control of EMT. Observed effects were obtained through simulation, starting from the basal Epithelial-like stable state and using an input configuration on the model that describes diverse EMT signals under RPTP activation conditions (IL6=ECM=EGF=HGF=DELTA=RPTP=1 and FAT4L=ROS=0).

| Type | Regulations | Targets | Model Perturbation | Simulation observations |
| --- | --- | --- | --- | --- |
| Single KO | Activation | p120 | p120[@RPTP0] | Multistability of phenotypes with Epithelial-like and partial EMT properties. |
| Single KO | Activation | Beta-catenin | bCat_mem[@RPTP0] | Phenotype with partial EMT properties. |
| Single KO | Inhibition | EGFR | EGFR[@RPTP0] | Phenotype with partial EMT properties. |
| Single KO | Inhibition | cMET | MET[@RPTP0] | Multistability with Mesenchymal-like and Partial EMT properties. |
| Double KO | Inhibition<br>Inhibition | EGFR<br>cMET | EGFR[@RPTP0]<br>MET[@RPTP0] | Multistability with Mesenchymal-like and Partial EMT properties. |
| Double KO | Activation<br>Activation | Beta-batenin<br>p120 | bCat_mem[@RPTP0]<br>p120[@RPTP0] | Multistability with Mesenchymal-like and Partial EMT properties. |
| Double KO | Inhibition<br>Activation | cMET<br>beta-catenin | MET[@RPTP0]<br>bCat_mem[@RPTP0] | Phenotype with partial EMT properties. |
| Triple KO | Inhibition<br>Activation<br>Activation | cMET<br>beta-catenin<br>p120 | MET[@RPTP0]<br>bCat_mem[@RPTP0]<br>p120[@RPTP0] | Mesenchymal-like phenotype. |
| Triple KO | Inhibition<br>Activation<br>Activation | EGFR<br>beta-catenin<br>p120 | EGFR[@RPTP0]<br>bCat_mem[@RPTP0]<br>p120[@RPTP0] | Phenotype with partial EMT properties. |

In addition, we have analyzed the activity of the nodes involved downstream of RPTP activation on the resulted model stable states and reconstructed a proposed mechanism of action for the control of EMT (Figure 3). The changes in activity of the nodes showed that the inhibition of cMET together with activation of p120 and β-catenin is critical to ensure the typical high Epithelial cell-cell adhesion. Unexpectedly, cMET inhibition by RPTP was necessary for preventing the inhibition of E-cadherin expression (CDH1) by SNAIL/ZEB via TCF/LEF activation of Wnt and TGFβ signalling. On the other hand, the combined inhibition of cMET with EGFR by RPTP was found to be required to ensure an absolute inhibition of the high focal adhesion dynamics via inhibition of FAK/SRC complex. Together, these results suggest that RPTPs need to target both catenins, MET and EGFR to prevent EMT and the stability of phenotypes with cell migration capacity.

**Figure 3.**
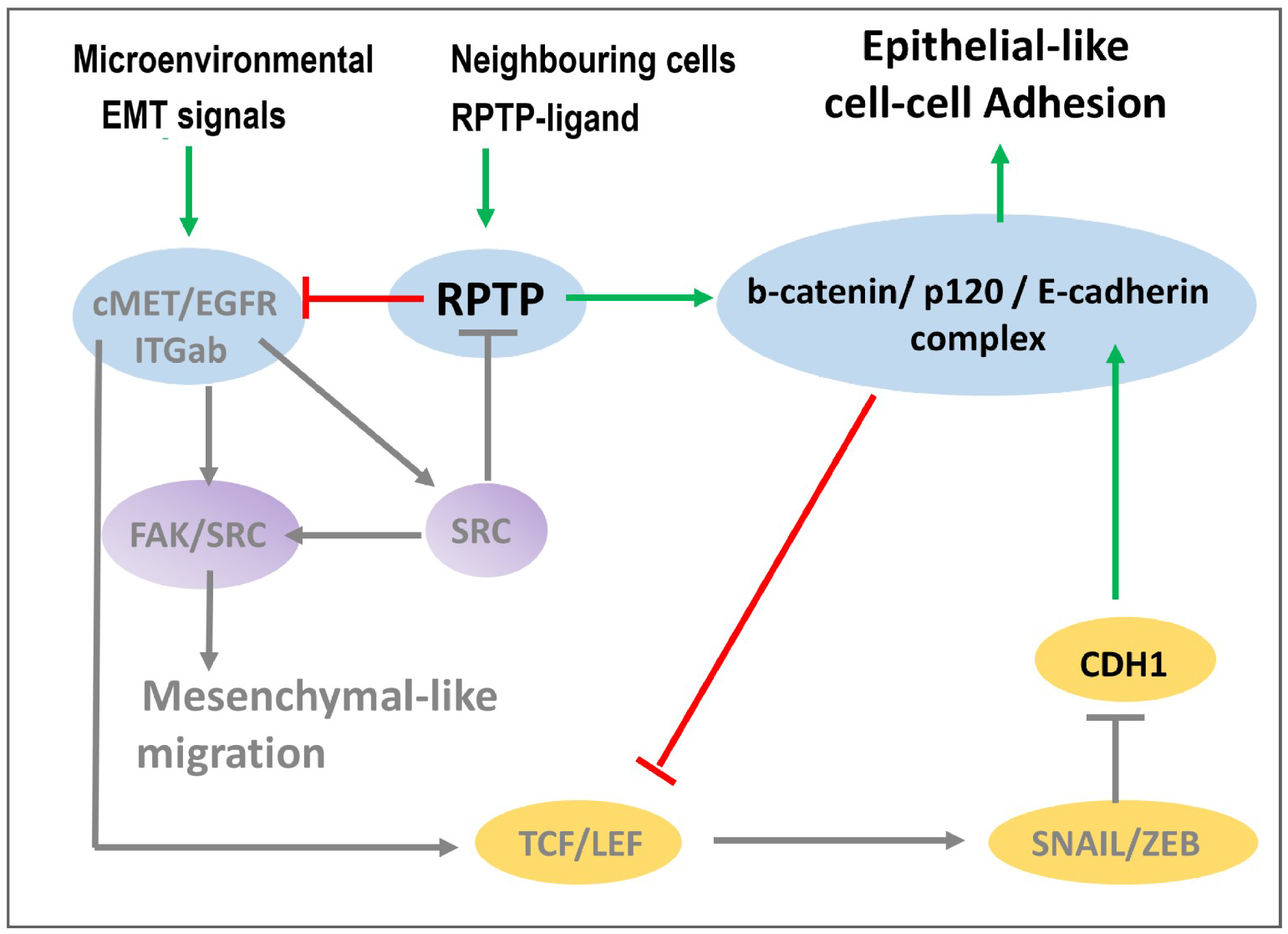
Proposed mechanism of EMT control by RPTP. Red arrows denote active inhibitions and green arrows denote active activations during simulations. Grey arrows indicate inactivity of processes during simulations. Active molecular components and phenotypic traits are indicated in black and inactive indicated in grey.

## 3. Discussion

In this work, we showed the potential capacity of RPTPs for controlling EMT by preventing EMT and promoting MET. This already makes an important contribution to the cancer field with the identification of a novel mechanism that could control cancer migration through EMT and eventually prevent metastasis [11,12,30]. In addition, it also provides a mechanistic explanation for the observed high cell contact induced MET in cancer cell lines [27,31]. In theory, high cell contact dependent RPTP signal is achieved in a scenario where a cell has enough neighbour cells expressing RPTP ligands, which would be the case of healthy epithelia [14]. On the other hand, low RPTP signal can be achieved in scenarios where enough neighbour cells are destroyed by either apoptosis or tissue damage such as wounds. This is compatible with our model results and conveys the idea that EMT is tightly controlled based on the demand for Mesenchymal cells, explaining the transient behaviour in wound healing [31].

Potentially, RPTPs can be a good inhibitor of EMT since they typically have higher activity rates (about 1000-fold higher) in comparison with RTK dependent growth factors signalling involved in EMT (e.g. EGFR and cMET) [15]. In general, RPTPs are highly expressed in Epithelial cells of most tissues, placing them as plausible candidates for a generic control mechanism [14]. However, only a restricted set of RPTPs have been proven to be controlled by cell contact interactions through homophilic or another type of ligand interactions [14]. In cancer, two cell contact dependent RPTPs, the RPTP-k and DEP-1, are often mutated or down-regulated [14,32]. This, together with the model analysis, supports the hypothesis that deregulations on RPTPs are relevant for cancer invasion and metastasis. In addition, RPTP-κ is reported to target both EGFR and β-catenin, whereas DEP-1 targets p120 and MET [14,33–35]. Based on the model analysis, all above mentioned targets were required for the EMT control by RPTPs. Thus, it is plausible to hypothesise that both RPTP-κ and DEP-1 may be collectively activated for effective control of EMT.

The model analysis further showed that oxidative stress generated during hypoxia plays a key role in inhibiting the control of EMT by RPTPs. This is particularly evident in RPTP-k of keratinocytes under UV, suggesting that it could also be the case of other sources of oxidative stress [28]. This places the antioxidant usage as a candidate for cancer therapy to prevent excessive accumulation of Mesenchymal-like cancer cells in tumours. Importantly, our analysis provides the conditions by which these Mesenchymal-like cancer cells may accumulate in tumours. However, not all conditions are plausible in the case of tumours growing in an epithelium, where cell contacts are high. This would exclude most analysed physiological scenarios based on the identified putative effect of FAT4, which may also prevent EMT but not trigger MET. Therefore, the tumour microenvironment under chronic inflammation conditions in combination with hypoxia is the most likely condition for promoting the accumulation of Mesenchymal-like cancer cells. According to model analysis, once these Mesenchymal-like cancer cells escape the primary tumour and migrate through blood, they can colonize other organs through MET if they encounter high cell contact and oxygen rich microenvironment. Thus, our results provide a mechanistic explanation for the correlation observed between metastasis and hypoxia under chronic inflammation conditions [8].

## 4. Conclusion

Based on the network model analysis showed here, cell contact dependent RPTP is demonstrated to be a potential control mechanism of EMT that can be tested in future experimental studies. Importantly, this control mechanism is shown here as a good hypothesis to explain MET by the microenvironment and explain the correlation between metastasis of carcinomas and hypoxia. Moreover, the analysis of the mechanism of RPTP control in this study provides candidate targets towards the design of new therapeutic strategies to prevent tumour cells to gain the capacity of invading the primary site.

## 8. Acknowledgements

The work presented in this paper was financially sponsored and hosted by BioenhancerSystems. FCT-Fundação para a Ciência e Tecnologia under the grant Ref SFRH/BD/52175/2013 for sponsoring the PhD thesis that resulted in the model used in this work. The Instituto Gulbenkian de Ciência that hosted and also financially sponsored the above mentioned PhD thesis. Dr Claudine Chaouiya for supervision and advice on model construction and validation. Eng. Pedro Fernandes for independent consulting on modelling biological systems.

## Abbreviations

(EMT): Epithelial-to-Mesenchymal Transition;
(MET): Epithelial-to-Mesenchymal-to-Epithelial Transition,
(ECM): extracellular matrix.

