## Supplemental data for "Simulation of multiple microenvironments shows a putative role of RPTPs on the control of Epithelial-to-Mesenchymal Transition"

### Supplementary data

#### 1. Initial stable states for simulations

The basal stable state of Epithelial-like and Mesenchymal-like phenotypes used as initial states during all simulations are presented in Figure S1. These stable states were obtained using simulation with all inputs (microenvironment signals) set to 0 and starting from all state space using the MonteCarlo algorithm implemented in GINsim. Only these two stable states (Figure S1) were reachable, resulting in one stable with cell adhesion properties that are characteristic of Mesenchymal-like and Epithelial-like phenotypes. A detailed discussion concerning the association of these stable states to these phenotypes phenotypes can be found in [1] .

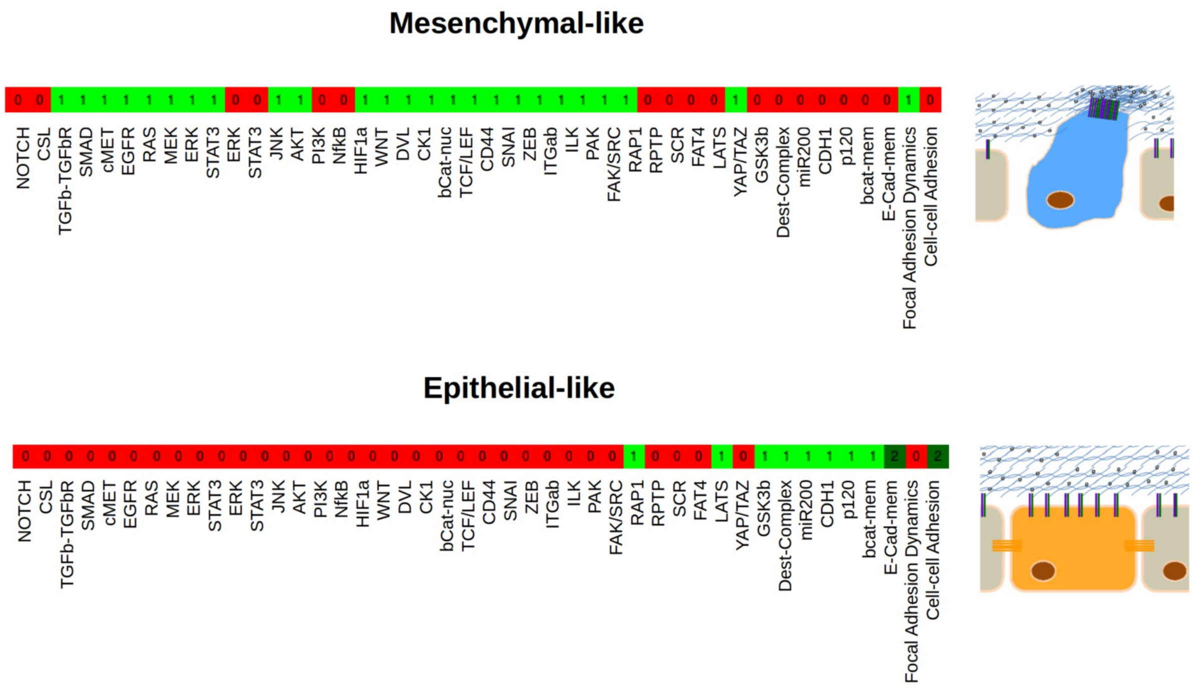


**Figure S1. Basal model stable states associated to Epithelial-like and Mesenchymal-like phenotypes.**

### 2. Model validation

The network model used for predictions was validated with phenotype and molecular observations from published experiments on epithelial-like cell lines described in Table S2. The observations were associated to discrete qualitative changes, which were used to compare with the simulations of these experiments. Full description on the methodology used and discussion is available in [2].

**Table S2: E**xperiments on mammalian epithelial cell lines used for model validation, grouped by perturbation type. The number of total observations (N) collected for each perturbation type.

| Perturbation | Experiments description | N | Refs. |
|---|---|---|---|
| **IL6+** | Exogenous addition of IL6 to normal mammary Epithelial cell lines. | 5 | [12] |
| **EGF+** | Exogenous addition of epidermal growth factor to several human Epithelial cell lines (cancer and normal). | 18 | [13–16] |
| **ECM+** | Exogenous addition of collagen type I or fibronectin to several human and mouse Epithelial cell lines (cancer and normal). | 10 | [16,17] |
| **EGF+ ECM+** | Exogenous addition of epidermal growth factor in combination with collagen type I or fibronectin to human breast, skin and squamous Epithelial cell lines (cancer and normal). | 5 | [16] |
| **HGF+** | Exogenous addition of hepatocyte growth factor to Epithelial cell lines from human, canine and mouse origins. | 35 | [5,14,18–20] |
| **WNT+** | Exogenous addition of Wnt protein to kidney canine and colon cancer Epithelial cell lines. | 8 | [5,21] |
| **TGFb+** | Exogenous addition of transforming growth factor beta to several human and mouse Epithelial cell lines. | 30 | [22–25] |
| **EGF+ ROS+** | Exogenous additions of epidermal growth factor with ROS intracellular generation in mammary Epithelial cell lines. | 10 | [26] |
| **P120 KO** | Knockout of p120 using siRNA in hamster ovary cell lines. | 8 | [3,4] |
| **B-Cat_mem KO** | Transformation of mouse embryonic and canine kidney Epithelial cell lines with a mutated form of E-cadherin incapable of binding to β-catenin. | 5 | [5,6] |
| **CK1 E1** | Overexpression of CK1 on human endothelial and canine kidney cell lines. | 7 | [7] |
| **CD44 E1** | Overexpression of CD44 in colon tumour cancer cell lines. | 5 | [8] |
| **SRC E1** | Overexpression of SRC in human breast and colon cancer cell lines. | 17 | [9–11] |

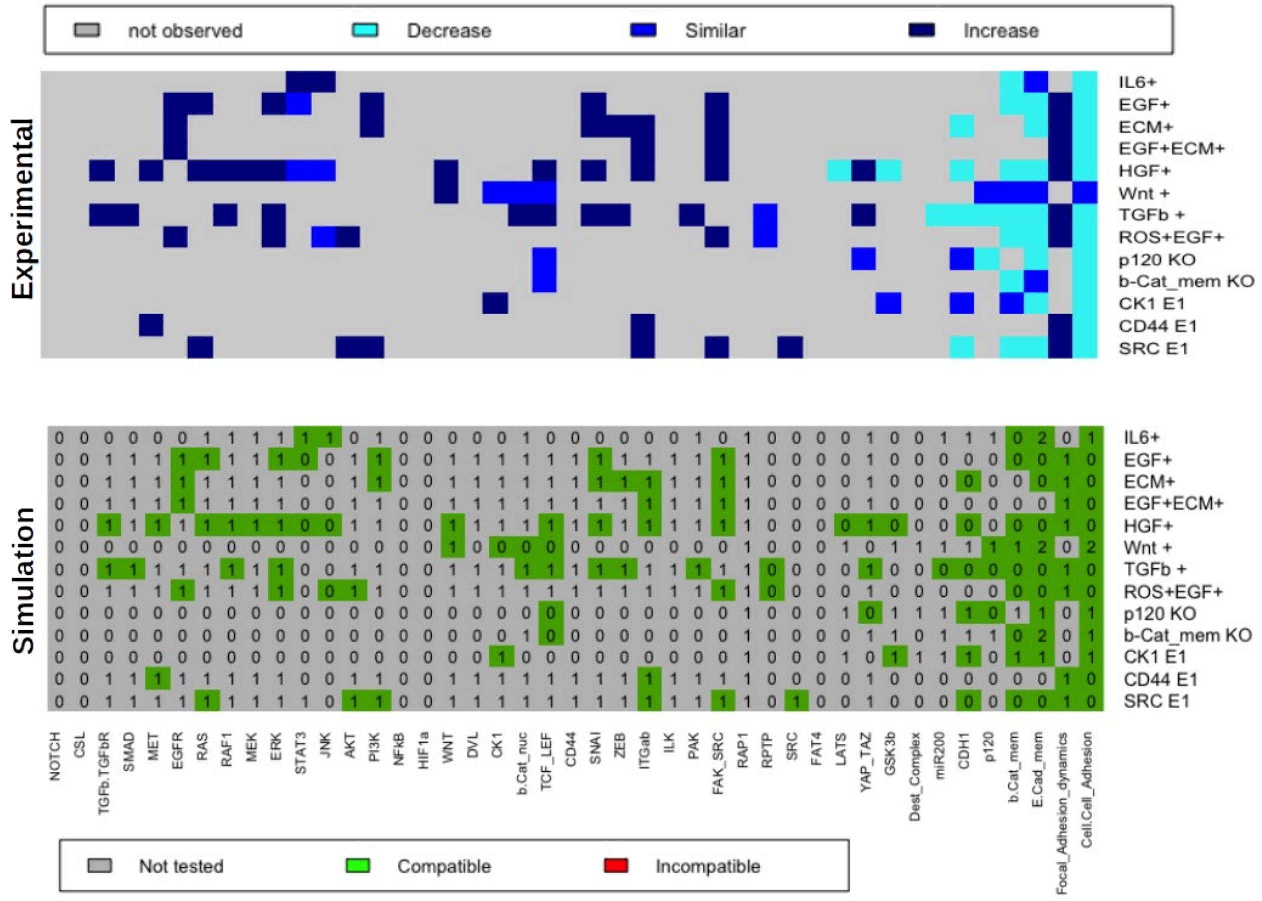


**Figure S1. Comparison of the observations of experiments on Epithelial cell lines (top) with simulation results (bottom).** Exogenous additions of microenvironment components are indicated by the symbol +, knockouts of intracellular network components by KO and overexpression/overactivation by the letter E. The value after the letter E indicate an ectopic activation at that value. For the experimental observations (top), the qualitative changes in activity are indicated by dark blue (increase), blue (no change) and cyan (decrease). For simulations (bottom), the value 0 means basal activity, value 1 intermediate (Multi-valued variables) or high activity (Boolean variables), and value 2 high activity (Multi-valued variables). Here, the nodes coloured with green indicate compatibility with experimental data and red incompatibility. Grey indicates predicted the activity of the nodes (not tested/observed experimentally).

### 3. Setup of physiological scenarios

For the simulation of physiological scenarios we define four well described physiological scenarios states which are summarized in table S2. For each of this four states we associate a correspondent model inputs configuration that better describes each of these states in a qualitative manner. Thus, for the scenarios that a given signal is reported to be highly abundant in the microenvironment we consider the value 1, otherwise we consider it to be 0.

**Table S2.** Model input variables used in simulations of particular physiological scenarios. For model inputs, the values 0 indicate basal and value 1 high degrees of activity.

| | Description | Model Inputs |
|---|---|---|
| **Healthy Epithelia** | Scenario where the Epithelial cells of an epithelia surrounding a tissue are not massively dividing under tissue repair or wound healing. In these conditions, the degree of growth factors, cytokines, ROS and ECM stiffness are low, considered here as basal (value 0) [27–29]. | IL6 = 0<br>ROS = 0<br>ECM=0<br>EGF=0<br>HGF=0 |
| **Tissue growth** | Scenario with high secretion of growth factors by Fibroblasts and adjacent Epithelial cells [27,30,31]. Stimulates cell proliferation to balance cell death. Occurs during tissue repair, tissue size homeostasis and in a tumour proliferative state. | IL6=0<br>ROS=0<br>ECM=0<br>EGF=1<br>HGF=1 |
| **Chronic Inflammation** | Scenario with a prolonged state of inflammation resulting in the accumulation of collagen I, IL6 cytokine and HGF secreted by recruited Fibroblasts and Macrophages to the inflammatory site [28,30]. Frequently observed during wound healing and in cancer. | IL6=1<br>ROS=0<br>ECM=1<br>EGF=0<br>HGF=1 |
| **Hypoxia** | Scenario achieved under fast growth of tumours, where oxygen available in the tumour microenvironment is consumed. Low levels of oxygen trigger the generation of high levels of intracellular ROS [32]. Frequently observed in invasive tumours in combination of chronic inflammation conditions [28,29,33]. | IL6=0<br>ROS=1<br>ECM=0<br>EGF=0<br>HGF=0 |
